# Structure of EPCR in a non-canonical conformation

**DOI:** 10.1101/2021.12.16.472967

**Authors:** Elena Erausquin, Adela Rodríguez-Fernández, Jacinto López-Sagaseta

## Abstract

Structural motion and conformational flexibility are often linked to biological functions of proteins. Whether the endothelial protein C receptor (EPCR), like other molecules, is vulnerable to folding transitions or might adopt alternative conformations remains unknown. The current understanding points to a rigid molecular structure suitable for binding of its ligands, like the anticoagulant protein C, or the CIDRα1 domains of *Plasmodium falciparum*. In this study, we have identified a novel conformation of EPCR, captured by X-ray diffraction analyses, whereby Tyr154 shows a dramatically altered structural arrangement, likely incompatible with protein C binding. Biolayer interferometry analysis confirms previous results supporting a critical role for this position in protein C binding. Importantly, the conformational change has no apparent effect in the bound lipid. We conclude these findings reveal a site of conformational vulnerability in EPCR and inform a highly malleable region that could modulate EPCR functions.

## Introduction

EPCR is an essential regulatory receptor that quickens the generation of the anticoagulant activated protein C (APC) on the surface of endothelial cells, which in turn prevents unhealthful levels of thrombin in blood. While EPCR mutations are linked to prothrombotic clinical outcomes^1^ and fetal death^2^ in humans, EPCR^-/-^ mice do not survive beyond the embryonic stage due to thrombotic associated lethality^3^, which mirrors the critical relevance of this receptor for proper development of life in mammals. Structurally, EPCR is a non-conventional MHC class I-like protein with a well-defined molecular architecture^4^. Like the CD1 family of receptors, EPCR presents bound lipids in a central and hydrophobic cavity.

A high-resolution crystal structure of the protein C Gla domain bound to EPCR^4^, and alanine mutagenesis studies^5^, identified the amino acids in EPCR proximal to the Gla domain that contribute and are essential for protein C/APC binding. Among these, Tyr154 plays a critical role as it establishes a nourished network of interactions with the Gla domain that likely sustains protein C/APC binding.

Through X-ray diffraction studies, we have identified a novel, non-canonical conformational state of EPCR with a strikingly unconventional fold in the Tyr154-Thr157 region. Particularly, a dramatic altered positioning of Tyr154 side chain is observed. This novel conformation in the α2 helix reveals a structurally vulnerable region in EPCR that could contribute to its varied biological functions.

## Results and discussion

Structural studies performed in our laboratory enabled crystallization of EPCR in an unusual conformation (Figures 1A-B and 2A). Structure solution readily showed F_o_-F_c_ difference signal indicating an evident conformational change in the α2 helix with particular impact in the orientation of Tyr154 side chain. Previous studies by Liaw and colleagues have demonstrated that Tyr154 side chain is essential for proper binding of protein C/APC to EPCR, as Tyr154 replacement with alanine results in an EPCR form unable to bind PC/APC^5^. This non-canonical rotamer of Tyr154 is the result of an alternative structural arrangement of a short loop that switches the direction of the α_2-1_ helix (Figures 1A-C). The hinge-like motif can be seen as a structural piece that breaks the α2 helix into two independent helical rigid bodies, α_2-1_ and α_2-2_. This is a common feature also observed in MHC class I and II antigen-presenting molecules^6^.

**Figure 1.**
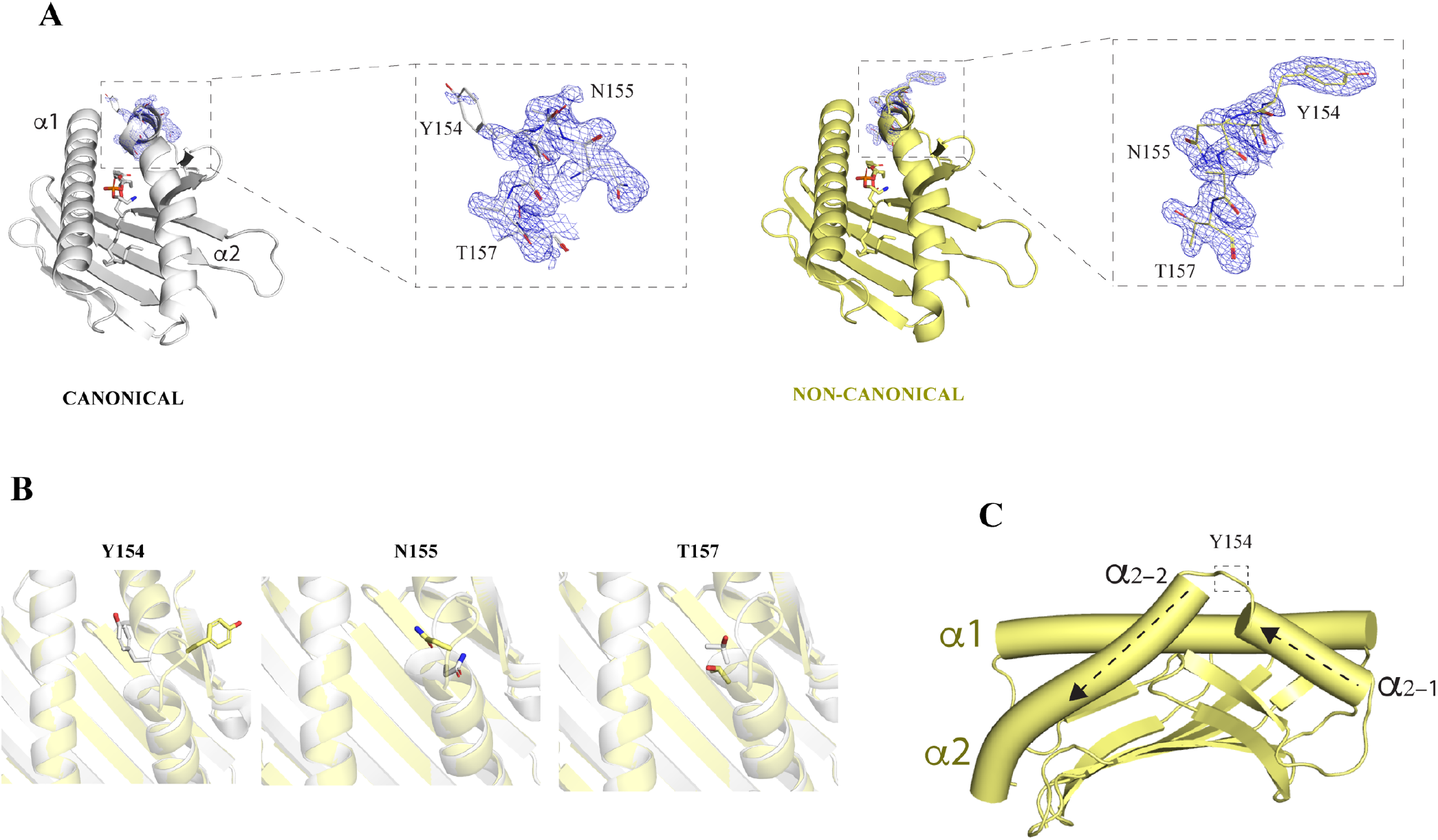
The non-canonical EPCR structure. (A) The canonical (left) and non-canonical EPCR structures are shown in grey and paleyellow colors, respectively. The residues in the α2 helix with severe folding transitions are highlighted. 2Fo-Fc electron density maps are shown as blue meshes. In order to confirm the novel conformation, maps were generated using either the canonical or non-canonical coordinates of residues Tyr154 through Thr157. The bound phospholipids are shown in the central cavities in stick format (B) Superposition of the canonical and non-canonical structures. Tyr154, Asn155 and Thr157 are highlighted for comparison purposes. (C) Directional switch in the α2 helix of EPCR.

Analysis of the interaction between EPCR and the protein C Gla domain shows how Tyr154 establishes numerous Van der Waals contacts with protein C Gla backbone and Asn2 and Phe4 side chains (figure 2A). A hydrogen-bond with Gla7 further contributes to the overall network of interactions mediated by Tyr154. In this novel structure, Tyr154 shows a profound structural transition and alters the location of its side chain in a manner such that is completely away from the protein C binding site. The protein backbone at this region also presents a deep rearrangement, starting with a rotation of the Ala153 carbonyl by 90 °, and followed by severe conformational shifts that affect not only Tyr154 but Asn155, Arg156 and Thr157 peptide bonds angles and side chains (Figure 1B). Arg156 is particularly flexible. While unliganded EPCR structures show a highly mobile Arg156 side chain, as inferred from the lack of electron density signals in previously reported structures, its position in the protein C Gla-bound structure is restricted by the interaction. This plasticity in the Arg156 side chain suggests that EPCR exists as a heterogeneous population whereby only those EPCR molecules with Arg156 in a favorable conformation enable protein C/APC binding. Or alternatively, this flexibility favors Arg156 motion and protein C docking. This is consistent in our structure, where we do not observe electron density signal for Arg156 side chain. From position 156, the receptor restores its canonical conformation at Arg158.

**Figure 2.**
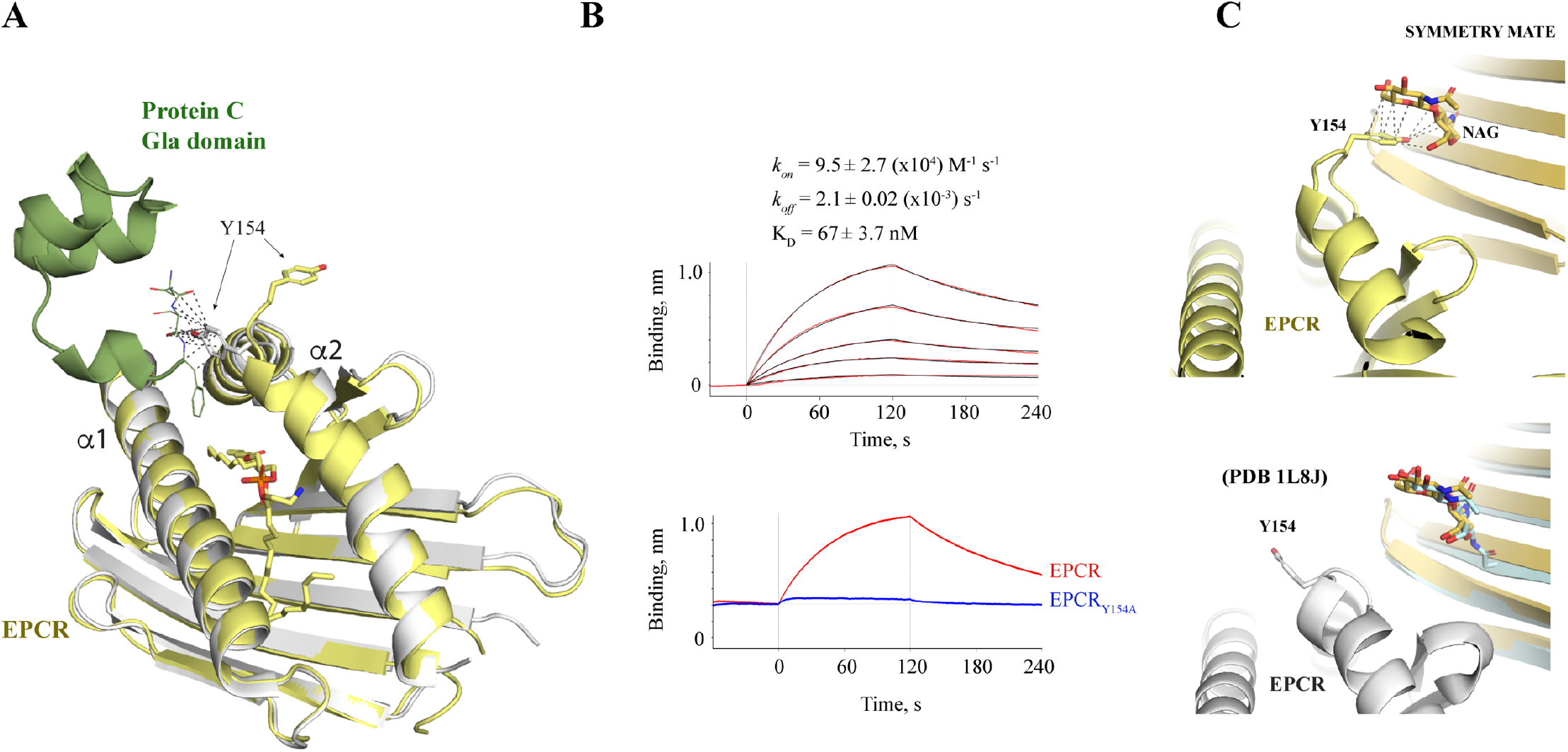
Impact of the non-canonical folding mode in protein C/APC binding. (A) Structure of the protein C Gla domain in complex with the canonical conformation of EPCR (PDB 1LQV). The contacts with the Gla domain established by EPCR Tyr154 are highlighted with grey dashed lines. Tyr154 residues in both the canonical and non-canonical EPCR structures are highlighted in sticks for comparison purposes and to better visualize the impact of the folding transition in protein C binding. (B) Upper panel, measurement of kinetic constant rates and affinity interaction between wild type EPCR and APC (upper panel). Red color traces denote buffer signal-substracted raw binding data and black traces indicate fitting to a 1:1 binding kinetic model. Lower panel, comparison of binding signal of 125 nM APC to EPCR or EPCR_Y154A_. (C) Upper panel, intermolecular contacts between the non-canonical EPCR Tyr154 and N-acetylglucosamine (NAG) in a crystallographic symmetry mate EPCR molecule. Lower panel, analogous view with the canonical EPCR structure (PDB 1L8J). The 1L8J symmetry mate (palecyan color) is shown superposed with the symmetry mate of the non-canonical EPCR structure.

In our crystal structure, each EPCR monomer have crystallographic symmetry mates with N-glycosylation molecules in the vicinity of Tyr154 (Figure 2C). It could be argued that this crystal packing and crystallographic contacts might have forced this novel conformation of the α_2-1_-α_2-2_ loop. However, our crystal recapitulates the space group and crystal packing of the previously EPCR structure (PDB 1L8J) solved by Oganessyan *et al*^4^. That is, the sugar molecules near to each EPCR monomer are also present in the previously determined structure, and therefore rules out a potential crystallization-induced conformation. Still, even if this alternative conformation was triggered by a close interacting molecule, it would reveal a site of vulnerability in a region of EPCR that is key for its anticoagulant properties.

To confirm the relevance of Tyr154 in protein C binding, we replaced it with alanine and monitored any potential binding to EPCR. As expected, and confirming previous findings by other groups^5,7^, replacement of EPCR Tyr154 with alanine has a profound impact in APC binding (Figure 2B). Consequently, our results support that Tyr154 is imperative for EPCR-mediated anticoagulation.

**Table 1.**
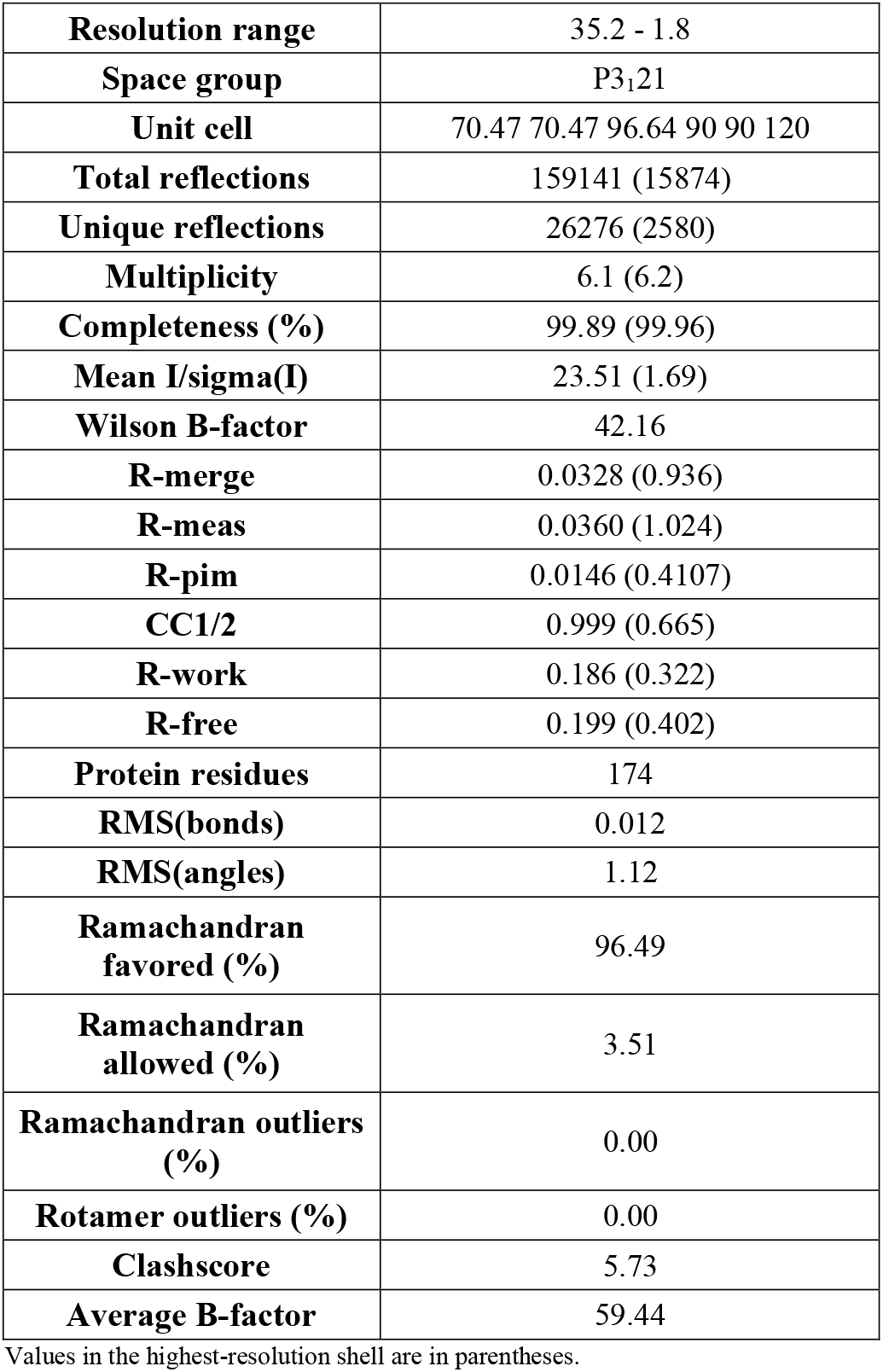
Diffraction data collection and refinement statistics.

The lipid ligand bound in the central cavity does not show any appararent alteration. As in the previous structure with the canonical EPCR structure, the electron density maps indicate the presence of a phospholipid molecule bound in the groove in a way similar to that already found by Oganesyan *et al*^4^.

Conformational heterogeneity and structural motion are inherent features often found in proteins. For instance, G protein-coupled receptors (GPCRs) possess structural motility that results essential for their biological properties^8,9^. Numerous crystallization studies have led to a deep understanding of GPCR molecular plasticity, which results in a wide spectrum of structural states. Another example of conformational plasticity is NFAT or the nuclear factor of activated T cells. X-ray structural studies of this transcription factor provide evidences for the coexistence of a heterogeneous population of fairly diverse conformations^10^.

In this study we have identified a novel conformation of EPCR that features a deep structural motion in Tyr154 and surrounding residues. This molecular transition associates with an EPCR state that lacks ability to bind protein C/APC. Our results point to a multivariate folding state of EPCR in physiological conditions. The structural plasticity of the hinge region represents a site of vulnerability that could be modulated by alternative binders. The question remains whether this novel folding arrangement represents a structural binding motif for other EPCR ligands, and which could determine relevant yet unknown roles of EPCR. In this line, recent works suggest EPCR-dependent T cell^11–13^ and antibody^14^ recruitment, which indicates that EPCR can interact with a broad variety of protein molecules.

In conclusion, this work reveals EPCR is not a receptor with a unique and rigid conformation “ready-to-bind” its ligands. EPCR presents a structurally vulnerable region in the α2 helix whose conformation may dictate EPCR properties in blood coagulation and recognition by immune receptors.

## Materials and Methods

### Recombinant production of EPCR

The extracellular region of human EPCR was produced in sf9 insect cells. Human EPCR cDNA (Genscript) was PCR amplified and cloned in frame with a GP64 signal peptide in a pAcGP67A transfer vector, using BamHI and NotI restriction enzymes and Optizyme™ T4 DNA ligase (Thermo Fisher Scientific). For crystallization purposes, the EPCR construct was prepared with an N-terminal 6xHis tag followed by a 3C protease cleavage site. For binding studies, we replaced this tag with a C-terminal 12xHis tag in a new EPCR construct. The Y154A substitution was prepared using the EPCR_12xHis_ template and complementary oligos containing the desired mutation. The first PCR products were used for a final overlapping PCR reaction that generated the final EPCR_Y145A_ construct with the C-terminal 12xHis motif. Sf9 insect cells (Gibco™) were transduced with the recombinant purified plasmids, BestBac 2.0 Δ v-cath/chiA Linearized Baculovirus DNA and Expres2 TR Transfection Reagent (Expression Systems) to produce recombinant baculovirus. All sequences were validated by Sanger sequencing (Stabvida).

### EPCR expression and purification

Sf9 insect cells were infected with EPCR amplified baculovirus, and the culture medium was collected after 72 hours of incubation at 28 °C in an orbital shaker. The culture supernatant was fred of cells by centrifugation and the clarified sample was concentrated using a 10 KDa MWCO Vivaflow (Sartorious) concentration device and dialysed overnight in HBS pH 7.4 buffer before purification with a Nickel NTA Agarose prepacked column (ABT). Protein was eluted with 20 mM Hepes 7.4 supplemented with 150 mM NaCl and 200 mM imidazole. Purified protein was digested overnight with in-house made 3C protease^13^ after imidazole removal. The tag-free protein was again loaded into the NiNTA Agarose cartridge and the flow through recovered. All purification steps were performed in an AKTA Pure 25M station (Cytiva). Protein purity was assessed by SDS-PAGE, then concentrated to 11.7 mg/mL for crystallization purposes.

### Crystallization and diffraction data collection

EPCR (11.7 mg/mL) was first screened against different crystallization reagents (Hampton Research and Molecular Dimensions) by the sitting drop vapour diffusion method. Initial hits were optimised in 24-well plates. The largest crystals appeared in 0.1 M sodium potassium tartrate, 20% PEG 3350 and recovered from the drop, soaked in the crystallization medium supplemented with 20% glycerol and cryo-cooled in liquid nitrogen prior to diffraction analyses.

### Structure determination and refinement

Diffraction data was collected at the Xaloc beamline, ALBA Synchrotron (Cerdanyola del Vallès). Data was indexed and integrated with XDS^15^, then scaled and merged with Aimless (CCP4 crystallographic suite)^16^. Previously deposited coordinates of EPCR (PDB accession number 1LQV) without ligands or water molecules were used as template to solve the structure using molecular replacement with Phaser^17^, which was then refined with phenix.refine^18^. Refinement strategies included XYZ, individual B-factors and optimization of X-ray/stereochemistry weight and X-ray/ADP weight. Initial steps were performed using rigid body refinement and Translation-Libration-Screw parameters were included in the final refinement processes. Conformational changes, ligands and water molecules were added guided by F_o_-F_C_ difference maps. The final molecule was generated after several cycles of manual building in Coot and refinement.

### C-terminal 12xHis EPCR expression and purification

Sf9 insect cells were infected with EPCR_WT_ or EPCR_Y154A_ baculovirus for 72 hours before culture medium collection. Culture supernatants were clarified and the proteins purified in HisGraviTrap columns (Cytiva). Proteins were eluted using 20 mM Hepes pH 7.4, 150 mM NaCl and 500 mM imidazole and buffer exchanged to 20 mM Tris pH 8.0 on a HiPrep 26/10 Desalting column (Cytiva) before further purification on HiTrap CaptoQ ImpRes IEX column (Cytiva). Pure protein was concentrated using 10 KDa Nanosep columns (Pall Corporation), aliquoted and frozen in liquid nitrogen for storage at −80°C.

### Biolayer interferometry

The impact of the Y154A substitution on EPCR was assessed by biolayer interferometry using the BLItz system (Sartorius). 12xHis-tagged EPCR_WT_ or EPCR_Y154A_ was immobilized on the surface of NiNTA pre-coated biosensors (Sartorius), at a concentration of 100 µg/mL, until stable levels were reached. The sensor was then pulsed with increasing concentrations of purified human APC (ThermoFisher) in 20 mM Hepes pH 7.4, 150 mM NaCl, 3 mM CaCl_2_ and 0.6 mM MgCl_2_. Interaction kinetics were calculated for each ligand and fitted to a 1:1 Langmuir binding model using the BLItz software (version 1.2.1.5).

## Acknowledgements

This work was supported by Ramón y Cajal (RYC-2017-21683) and Generación de Conocimiento (PGC2018-094894-B-I00) grants to JLS from the Ministry of Science and Innovation, Government of Spain. We thank the staff of XALOC beamline at ALBA Synchrotron for their assistance with X-ray diffraction data collection. We also thank Maria Gilda Dichiara Rodríguez for her technical support throughout this work.

## Data availability

Coordinates and structure factors have been deposited in the Protein Data Bank and have been assigned the accession code 7Q5D.

## Authorship contributions

Conceptualization: JLS; Experimental procedures: JLS, EEA and ARF; Data analysis: JLS, EEA and ARF; Manuscript writing: JLS, EEA.

## References

1. Dennis, J. et al. The endothelial protein C receptor (PROCR) Ser219Gly variant and risk of common thrombotic disorders: a HuGE review and meta-analysis of evidence from observational studies. Blood 119, 2392–2400 (2012).

2. Franchi, F. et al. Mutations in the thrombomodulin and endothelial protein C receptor genes in women with late fetal loss. Br. J. Haematol. 114, 641–646 (2001).

3. Gu, J.-M. et al. Disruption of the endothelial cell protein C receptor gene in mice causes placental thrombosis and early embryonic lethality. J. Biol. Chem. 277, 43335–43343 (2002).

4. Oganesyan, V. et al. The crystal structure of the endothelial protein C receptor and a bound phospholipid. J Biol Chem 277, 24851–24854 (2002).

5. Liaw, P. C., Mather, T., Oganesyan, N., Ferrell, G. L. & Esmon, C. T. Identification of the protein C/activated protein C binding sites on the endothelial cell protein C receptor. Implications for a novel mode of ligand recognition by a major histocompatibility complex class 1-type receptor. J. Biol. Chem. 276, 8364–8370 (2001).

6. Zacharias, M. & Springer, S. Conformational flexibility of the MHC class I alpha1-alpha2 domain in peptide bound and free states: a molecular dynamics simulation study. Biophys. J. 87, 2203–2214 (2004).

7. Sampath, S. et al. Plasmodium falciparum adhesion domains linked to severe malaria differ in blockade of endothelial protein C receptor. Cell. Microbiol. 17, 1868–1882 (2015).

8. Mary, S. et al. Ligands and signaling proteins govern the conformational landscape explored by a G protein-coupled receptor. Proc. Natl. Acad. Sci. U. S. A. 109, 8304–8309 (2012).

9. Luttrell, L. M. & Kenakin, T. P. Refining efficacy: allosterism and bias in G protein-coupled receptor signaling. Methods Mol. Biol. 756, 3–35 (2011).

10. Stroud, J. C. & Chen, L. Structure of NFAT bound to DNA as a monomer. J. Mol. Biol. 334, 1009–1022 (2003).

11. Willcox, C. R. et al. Cytomegalovirus and tumor stress surveillance by binding of a human gammadelta T cell antigen receptor to endothelial protein C receptor. Nat. Immunol. 13, 872–879 (2012).

12. Mantri, C. K. & St John, A. L. Immune synapses between mast cells and gammadelta T cells limit viral infection. J. Clin. Invest. (2018). doi:10.1172/JCI122530

13. Erausquin, E. et al. A novel α/β T-cell subpopulation defined by recognition of EPCR. bioRxiv 2021.07.01.450412 (2021). doi:10.1101/2021.07.01.450412

14. Müller-Calleja, N. et al. Lipid presentation by the protein C receptor links coagulation with autoimmunity. Science 371, (2021).

15. Kabsch, W. XDS. Acta Crystallogr D Biol Crystallogr 66, 125–132 (2010).

16. Evans, P. R. & Murshudov, G. N. How good are my data and what is the resolution? Acta Crystallogr. D. Biol. Crystallogr. 69, 1204–1214 (2013).

17. McCoy, A. J. et al. Phaser crystallographic software. J. Appl. Crystallogr. 40, 658–674 (2007).

18. Adams, P. D. et al. PHENIX: a comprehensive Python-based system for macromolecular structure solution. Acta Crystallogr D Biol Crystallogr 66, 213–221 (2010).

